# Duplex Proximity Sequencing (Pro-Seq): A method to improve DNA sequencing accuracy without the cost of molecular barcoding redundancy

**DOI:** 10.1101/163444

**Authors:** Joel Pel, Wendy W. Y. Choi, Amy O. Leung, Gosuke Shibahara, Laura Gelinas, Milenko Despotovic, W. Lloyd Ung, Andre Marziali

**Affiliations:** Boreal Genomics Inc.; Department of Physics and Astronomy, University of British Columbia

**Keywords:** Next generation sequencing, Liquid biopsy, Barcoded sequencing, Molecular barcoding, Duplex sequencing, Rare variants, Unique molecular identifiers (UMI), Cell-free DNA (cfDNA), Circulating tumor DNA (ctDNA), Formalin Fixed Paraffin Embedded (FFPE) Tissue

## Abstract

A challenge in the clinical adoption of cell-free DNA (cfDNA) liquid biopsies for cancer care is their high cost compared to potential reimbursement. The most common approach used in liquid biopsies to achieve high specificity detection of circulating tumor DNA (ctDNA) among a large background of normal cfDNA is to attach molecular barcodes to each DNA template, amplify it, and then sequence it many times to reach a low-error consensus. In applications where the highest possible specificity is required, error rate can be lowered further by independently detecting the sequences of both strands of the starting cfDNA. While effective in error reduction, the additional sequencing redundancy required by such barcoding methods can increase the cost of sequencing up to 100-fold over standard next-generation sequencing (NGS) of equivalent depth.

We present a novel library construction and analysis method for NGS that achieves comparable performance to the best barcoding methods, but without the increase in sequencing and subsequent sequencing cost. Named Proximity-Sequencing (Pro-Seq), the method merges multiple copies of each template into a single sequencing read by physically linking the molecular copies so they seed a single sequencing cluster. Since multiple DNA copies of the same template are compared for consensus within the same cluster, sequencing accuracy is improved without the use of redundant reads. Additionally, it is possible to represent both senses of the starting duplex in a single cluster. The resulting workflow is simple, and can be completed by a single technician in a work day with minimal hands on time.

Using both cfDNA and cell line DNA, we report the average per-mutation detection threshold and per-base analytical specificity to be 0.003% and >99.9997% respectively, demonstrating that Pro-Seq is among the highest performing liquid biopsy technologies in terms of both sensitivity and specificity, but with greatly reduced sequencing costs compared to existing methods of comparable accuracy.

## Introduction

The ability to detect rare DNA variants in a background of healthy DNA using next generation sequencing (NGS) has enormous potential to impact diagnostics in oncology, and prenatal testing. In cancer diagnostics, the detection of circulating tumor DNA (ctDNA) among cell-free DNA (cfDNA) in peripheral blood has enabled non-invasive detection and profiling of many types of cancers [1-4]. These “liquid biopsies” have been shown to provide actionable information in a significant fraction of patient cases [1, 4].

Initially, the promise of liquid biopsies was limited technically by the relatively high error rate of NGS systems, as true ctDNA mutations were obscured by inherent errors in DNA library preparation and sequencing. Modern NGS systems typically produce errors at a per-base rate of 10^-2^ to 10^-3^ [5-7], while clinically relevant mutations have been shown to be at or below that level [1, 8], making many true variants undetectable. A number of barcode-based (or UMI-based: Unique Molecular Identifier) error correction strategies have been developed in recent years [4, 9-23] but most of these methods increase the amount of sequencing required per sample. As the technical challenges of liquid biopsy assays are overcome, a major challenge remaining for broad clinical adoption of liquid biopsies is the increased cost associated with sequencing redundancy per sample [1]. Additionally, implementation of error correction has increased assay complexity and workflow time, to multiple days in many cases, introducing additional logistical barriers to clinical adoption.

In general, barcoding methods work by uniquely labeling (barcoding) a starting nucleic acid molecule (either by ligation or PCR), targeting the analysis to a specific genomic region of interest through target capture or further PCR, and then making redundant PCR copies of each target (Fig 1A). The amplified pool of redundant copies is sequenced, after which reads are grouped *in silico* into “families” based on their unique labels. Since each label represents a unique starting molecule, a consensus sequence can be determined for each read family, assuming sufficient copies are present. The typical average number of copies, or reads, per family required to make a consensus is around 20 [18, 24], which represents the fold-increase in sequencing required to achieve low error rate. For example, if a sequencing depth (or coverage) of 10,000 unique targets or genomes is desired for low frequency mutation detection, a total of 200,000 fold ‘depth’ is required when barcoding redundancy is included. Combining barcoding with *in silico* ‘polishing’, these techniques can reduce the per-base error rate to 10^-5^ errors per base [18].

**Fig 1.**
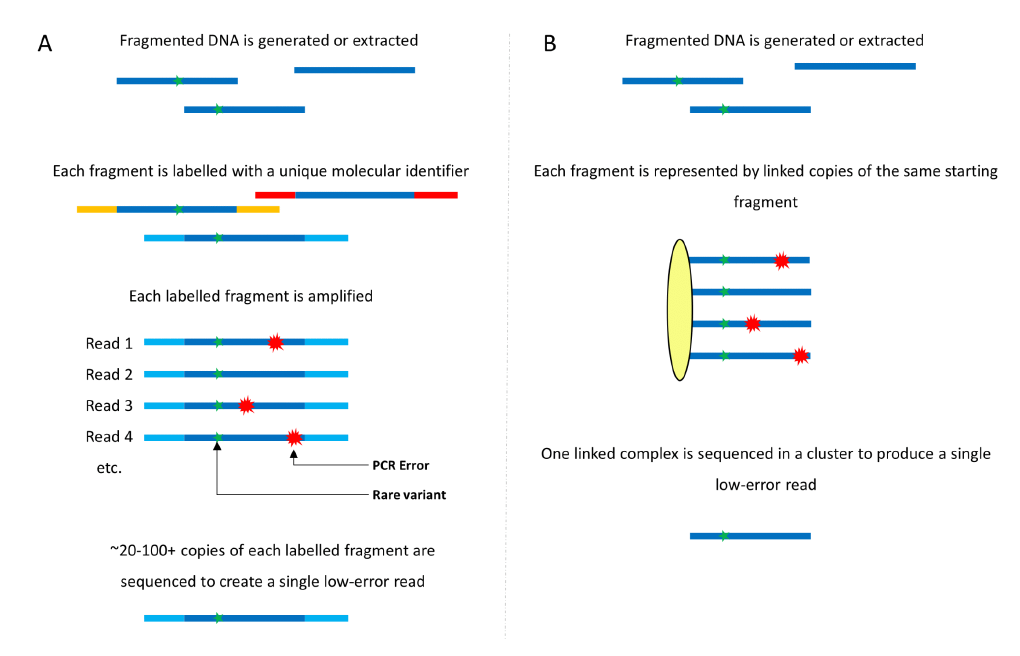
Barcoding vs. Pro-Seq. (A) Common molecular barcoding/UMI methods involve uniquely labeling each DNA molecule with a molecular identifier or barcode. Many copies of each barcoded molecule are sequenced, and reads from individual fragments are collected in software. True variants should be common to every read, while errors should only occur in a smaller fraction of the reads. 20 or more reads are often required to generate a software consensus for a single low-error read. (B) Pro-Seq physically links copies of the same starting fragment into a single complex. Each complex is then sequenced in a single cluster, producing a high-fidelity read without redundancy.

Further reduction in error rate has been achieved through a method called ‘duplex sequencing’ [11]. This method is similar to the barcoding scheme described above, except that starting molecules are labeled with barcodes through ligation in such a way that both senses of the starting molecule can be collapsed into a single barcode family, requiring true variants to be present on both senses of the starting duplex. Duplex sequencing has been shown to reduce errors to below 10^-6^ errors per base [25] and has the powerful ability to detect and reject DNA damage and rare sources of errors such as “jackpot mutations” (errors in the first cycle of PCR), which are not generally corrected in single-stranded barcoding. This is especially useful when working with potentially damaged DNA such as FFPE [26], or looking for very rare mutations in early cancer detection [1]. This performance comes at a cost however, as the average duplex family size can be greater than 100 [25], which correlates to a 100 fold-increase in the number of sequencing reads required per sample compared to regular NGS. Additionally, barcoding methods typically suffer from increased PCR bias and workflow complexities due to the presence of barcodes [13], further limiting clinical deployment. Several whole-genome barcoding methods also exist [27, 28], but remain exceptionally expensive unless coverage is very low (∼1x).

At least one technology has attempted to reduce the sequencing required for barcoding methods while retaining low error rate. Circle sequencing [13] uses a rolling circle approach to make concatenated copies of each starting molecule that can be read in single sequencer cluster. Correcting for DNA damage by chemical means, they have demonstrated per-base error rates down to ∼10^-6^. While sequencing usage is reduced compared to conventional barcoding, there are still several limitations of this method.

A practical limitation is the read length required to read more than two template copies in the concatenated template structure, which limits the error rate achievable. Since cell free DNA (cfDNA) is on average ∼170bp [29], it is only practically possible to read the single copies on each end of the concatenated template with a paired-end sequencing strategy, such as is available on Illumina platforms. Also, long concatenated templates are known by the manufacturer to inhibit cluster generation, reducing usable sequencing clusters. Additionally, in its current form, the technique is not able to create concatenated duplex reads, thus requiring extra sequencing if duplex information is desired.

We have developed Proximity Sequencing (Pro-Seq), a library preparation method that solves these challenges by physically merging both senses of read families into a single cluster and using the sequencer to generate a family consensus, thus eliminating the use of barcodes and redundant reads (Fig 1B). Here we describe the Pro-Seq method, report the analytical characterization of the assay and demonstrate its utility for high accuracy liquid biopsy with significantly reduced sequencing requirements, and a simple, one day workflow.

## Proximity Sequencing (Pro-Seq) Method

The Pro-Seq method is illustrated for an Illumina® sequencer in Fig 1B and Fig 2, and is conceptually applicable to other sequencing-by-synthesis platforms as well. In its general form, the method involves linking multiple copies of a single DNA template at the 5’ end early in the workflow so that the sequences of all molecules in a linked complex are nominally the same, with the exception of any errors made in their derivation from the parent strand. The linking is arranged in such a way that both senses of the starting template can be represented in a single linked complex, providing duplex information.

**Fig 2.**
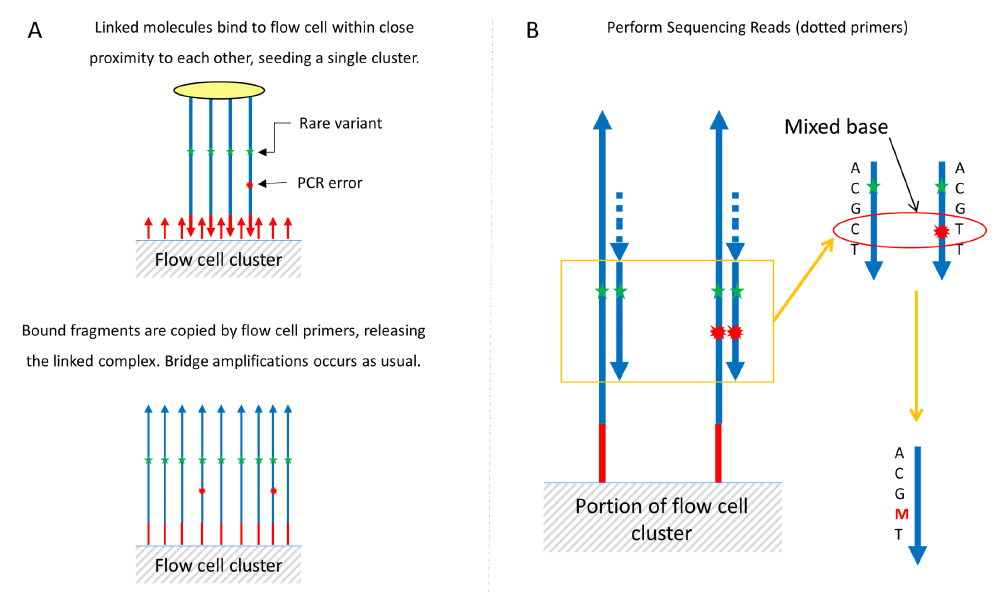
Pro-Seq sequencing. (A) Linked molecules are bound to the flow cell in close proximity to each other and form a single cluster as the size scale of the linker is much smaller than the size of a cluster. Following standard cluster generation, the bound fragments are extended and the linked template washed off (they do not interfere with flow cell function). After extension, bridge amplification proceeds as normal, with each cluster represented by multiple copies of the same starting molecule. (B) Clusters are sequenced, automatically generating an average or consensus of each base position, eliminating errors that occur as a small fraction of a cluster. In the case where the error signal is of similar scale to the true signal, error positions can be identified as mixed bases and masked (‘M’).

The linked complex is then sequenced directly so that the multiple linked copies seed a single sequencing cluster/colony (Fig 2). Cluster generation proceeds as usual, except that a single cluster now represents the aggregation of multiple redundant members of a family, instead of a single molecule. As sequencing proceeds, errors that are low abundance within an individual cluster are suppressed automatically by the sequencer’s basecaller. After sequencing, additional error bases are identified *in silico* by a drop in relative fluorescence (fQ), and subsequently masked (Fig 3). The outcome is a collapsing of multiple reads from a single starting template into a single cluster, increasing the accuracy of each cluster on the sequencer rather than requiring many clusters to achieve the same result. Depending on the application, it is also possible to integrate unique molecular identifiers for counting purposes, ensuring accurate quantification of sequenced molecules. We have developed both targeted and whole genome workflows based on this concept, but the targeted approach is the focus of this manuscript. Whole Genome Pro-Seq is described in S1 Fig.

**Fig 3.**
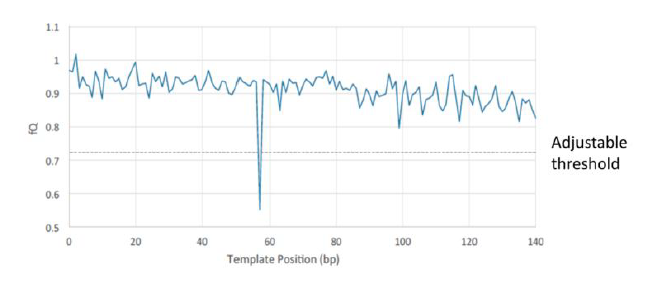
Pro-Seq error identification. In many cases, errors are corrected automatically on the sequencer as they represent a minority sequence compared to the dominant base within a cluster, and are ignored or not detected by the basecaller. To check for errors (mixed bases) that are of similar frequency to the correct base, the relative fluorescence (fQ) is calculated for each base in a read, in such a way that dips represent the presence of a mixed base. An adjustable threshold is used to identify dips and only the mixed base position is then masked. The rest of the read can be trusted to provide high-fidelity sequence information.

The targeted Pro-Seq workflow is outlined in Fig 4, and described in detail in the Materials and Methods. Briefly, the simple workflow consists of three main steps: droplet PCR, enzymatic cleanup and sequencing. Non-denatured dsDNA is loaded directly into droplets to retain duplex information, at a concentration that yields on average zero or one target template contained in each drop (ssDNA can also be sequenced in the same way with low error rate, but will not benefit from duplex error correction). Each droplet contains all multiplex gene specific primer sets, as well as universal linked primers with sequencing adapters. After droplets are loaded, the PCR reaction is thermally cycled to create linked molecules from each template-containing drop (effectively performing gene specific and universal PCR simultaneously). The emulsion is then broken and un-linked DNA is digested so only linked DNA remains. After quantification, the library is sequenced. The workflow is rapid, as a single technician can easily process multiple samples from extracted DNA to loaded sequencer in less than an 8-hour work day.All data presented in this paper uses a primer linking two molecules; however, constructs with up to 100 linkers have also been tested. These higher order linkers may reduce error rate further than what is reported herein.

**Fig 4.**
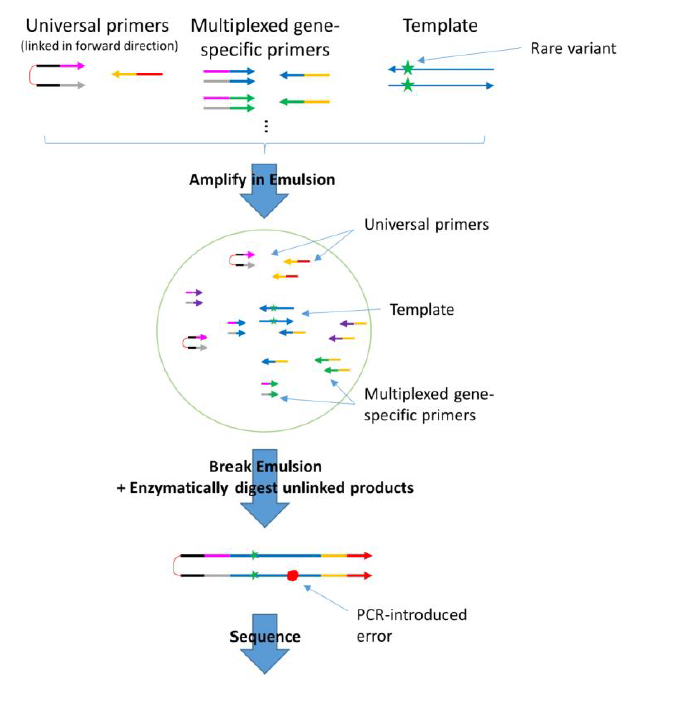
Overview of the targeted Pro-Seq workflow. (described in detail in the Materials and Methods). In brief, double stranded DNA is loaded directly into droplets such that on average zero or one template molecule is incorporated in each droplet. Off-target DNA (not shown in figure) is also loaded into droplets, but does not amplify. Within each droplet are multiplexed gene-specific primers, and the Pro-Seq universal 5’ PEG-linked primers. The droplets are PCR cycled such that all copies of the starting template are linked to the universal linked primers (shown in detail in S2 Fig). The emulsions are then broken, and the un-linked strands are digested and cleaned up. After quantification, the library is ready for sequencing.

## Results

We sought to evaluate and compare the analytical specifications of Pro-Seq to existing methods in order to assess its suitability for liquid biopsy applications, as many groups have previously shown the clinical utility of liquid biopsy for given assay characteristics [1, 2, 4, 30]. In addition, as a secondary result, we characterized the background mutation frequency in cell line cfDNA standards, demonstrating that care must be taken when using this source of DNA as a standard in high sensitivity assays.

Analytical specificity (or analytical true negative rate) is defined as the fraction of truly negative samples that are called negative. It can also be defined as 1 – FPR, where FPR is the False Positive Rate and in our case is defined per sample as the total number of non-reference bases called (regardless of abundance) divided by total bases called. This metric was used to provide an absolute measure of assay performance (per base), and, notably, is different than many other assay performance reports which define false positive rate as the rate of inadvertently calling a mutation above a certain threshold frequency [4, 18].

The targeted Pro-Seq false positive rate (FPR) was measured using a 7-amplicon panel on wild-type plasma-derived cell-free DNA (IPLAS – K2 EDTA, Innovative Research, Novi, MI), and was found to be 2.6 x 10^-6^ errors per base (n = 12, SD = 1.1 x 10^-6^). As a reference for a larger panel, the FPR for a 19-plex Pro-Seq assay was measured to be 1.1 x 10^-6^ errors per base (Fig 5). Only Pro-Seq error correction was used in analysis; no ‘polishing’ [18] or other *in silico* error reduction methods were employed, which we expect would lower the FPR further. The 7-plex FPR results in a per-base analytical specificity of 99.9997%.

**Fig 5.**
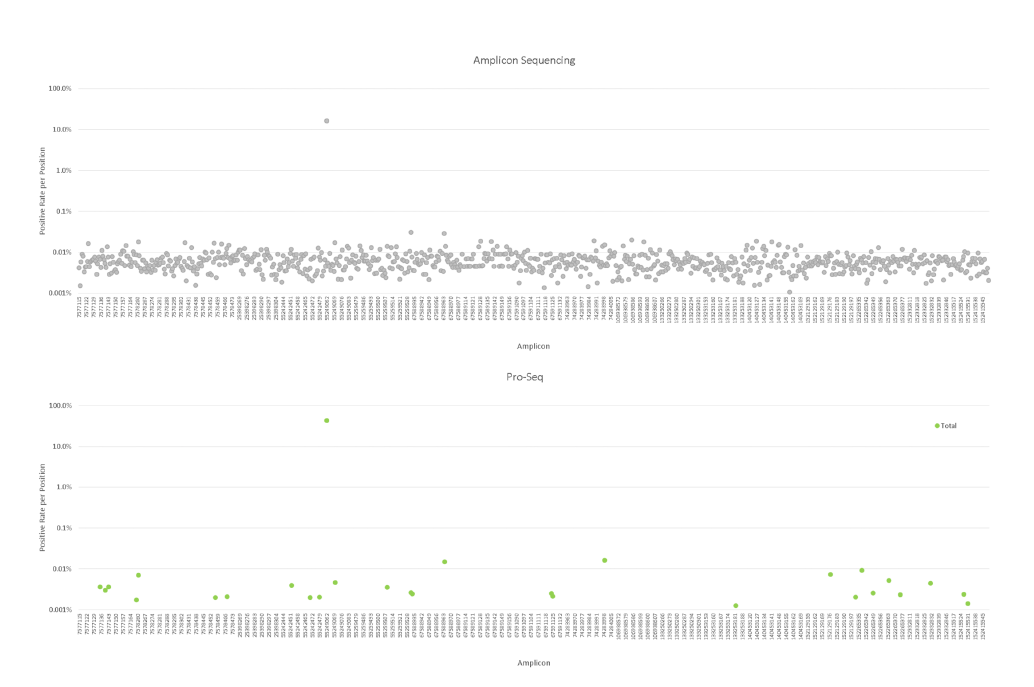
Comparative sequencing performance between amplicon sequencing with high fidelity polymerase (top) and Pro-Seq (bottom). A wild-type plasma sample was sequenced using a 19-amplicon panel with both methods, and the FPR plotted per base position. Amplicon sequencing (grey) has an average FPR of 1.2 x 10^-4^ errors per base, compared to Pro-Seq (green), which had an average FPR of 1.1 x 10^-6^ errors per base (a known SNP at panel position 237 is ignored for FPR calculations).

Analytical sensitivity (or analytical true positive rate) is defined as the fraction of truly positive mutations that are detected as positive. We characterized this sensitivity in two ways. First, by measuring the molecular sensitivity, where we fixed the number of input genomes and measured our ability to detect SNV or indel-containing molecules as a function of the number of mutant copies. This was done by titrating replicates of cell line DNA with five known mutations at specific positions into plasma-derived DNA from a healthy donor, known to be wild-type at the same positions (see Materials and Methods). Positive mutation detection was set to be above a threshold of 0.5 genomes, and the resulting data is presented in Table 1. In the lowest abundance sample, containing an average of 1.5 copies of each mutant, mutant copies were detected successfully for over 70% of the theoretically accessible mutations as estimated by sampling statistics. This increased to 100% detection between 4.5 and 15 copies per mutant.

**Table 1.** Molecular Sensitivity Characterization.

| Expected Number of<br>Copies per Mutant | Expected Number<br>of Mutants<br>(both replicates) | Sampling-corrected<br>Number of Mutants | Total Number of<br>Mutants Detected | Fraction of<br>Mutants Detected<br>(sampling corrected) |
| --- | --- | --- | --- | --- |
| 45 | 10 | 10.0 | 10 | 100% |
| 15 | 10 | 10.0 | 10 | 100% |
| 4.5 | 10 | 9.9 | 7 | 71% |
| 1.5 | 10 | 7.8 | 6 | 77% |
| 0 | 0 | 0.0 | 0 | 0% |

Second, we characterized analytical sensitivity by the detection threshold, using a metric defined in [4] as the SNV fraction at which ≥80% of SNVs were detected above wild-type background. We did this by fixing the number of SNV molecules at ten, above the molecular sensitivity and sampling limits, and then by increasing the number of wild-type genomes to reduce the variant fraction. Cell line DNA carrying the same five known mutants as presented above was titrated in duplicate into increasing amounts of wild-type cell line DNA, to generate samples with the desired mutant fractions. Wild-type cell line DNA with no mutant spike was also analyzed to measure background mutation levels. The detection threshold was measured to be 0.003%, as the lowest mutant fraction with four of five mutants detected above background. 100% of mutations were detected at 0.01% mutant fraction. Wild-type cell line samples analyzed at the same depth as the 0.003% replicates showed positive background detection for EGFR T790M, but the other four mutants showed no background. It is important to note, especially in the case of cell line DNA, that the EGFR mutation detected in the wild-type sample may be a real variant. The average expected vs. average measured frequency across the five mutations is shown in Fig 6, and is concordant across the tested range.

**Fig 6.**
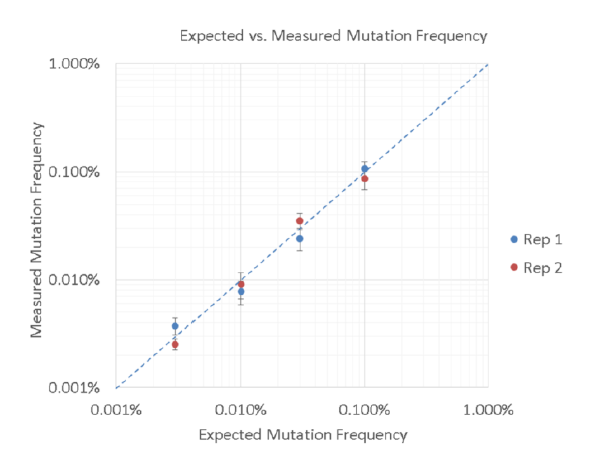
Expected vs. Measured Mutation Frequency. The average measured mutation frequency across all five mutations is plotted against the average expected frequency, for two replicates. Error bars indicate the standard error of the mean at each point, while the dashed line indicates 1:1 concordance between expected and measured values. Data is shown in S6 Table.

To assess the impact of duplex information in Pro-Seq we measured the prevalence of G>T (‘G-to-T’) and subsequent C>A variants, compared to the other ten variant possibilities, for the same 12 wild-type plasma runs used to assess FPR. G>T transversions are often associated with DNA damage from sample handling or library prep [26, 31], leading to higher representation of G>T and subsequent C>A variants compared to other variants in the absence of duplex correction. Additionally, ‘jackpot mutations’, i.e. errors that happen very early in PCR, may introduce a sequence and strand specific bias for certain mutation types, if not corrected.

The data presented in Fig 7 demonstrate comparable G>T and C>A frequency compared to common errors C>T and G>A [18, 31], suggesting damage or other errors occurring early in Pro-Seq do not dominate the false positive rate. Also noteworthy is the fact that complementary mutation types are well balanced (ex: A>G and T>C), suggesting that both strands of the starting duplex are evenly represented [11]. The observed discrepancy between A>C and T>G mutation rates may be explained by sampling noise, since these two mutation types typically occurred zero or once during each run, possibly leading to inaccurate measurements due to few data points.

**Fig 7.**
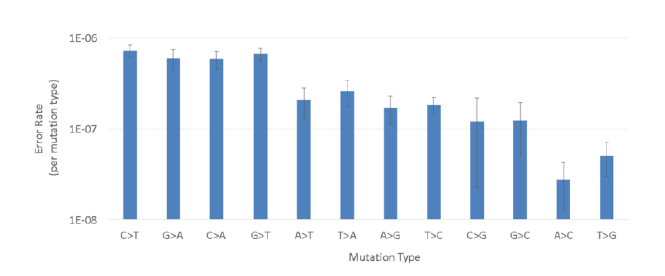
Average mutation rate as a function of mutation type across the 12 runs used to measure FPR. The error rate was calculated per run as the count of all non-reference calls per mutation type over total bases sequenced. Error bars represent the standard error of the mean.

Pro-Seq was also characterized by how efficiently it uses the sequencer, compared to other methods. Since barcoded sequencing methods typically report the number of reads required to make a consensus for each individual input template molecule, we sought to compare Pro-Seq by this metric. Though Pro-Seq does not use consensus reads, there is a fraction of reads that are not seeded by two or more templates, and thus a measurement of the number of reads required to generate a single high fidelity read is still appropriate for comparison. Sequencing efficiency was characterized by measuring the average number of on-target reads required to achieve a single high fidelity read, as a function of the measured cfDNA error per base. This measurement was made using the workflow described in Materials and Methods, and the data is presented in Fig 8, along with estimates made for other methods. Fewer reads per consensus corresponds proportionally to reduced sequencing cost.

**Fig 8.**
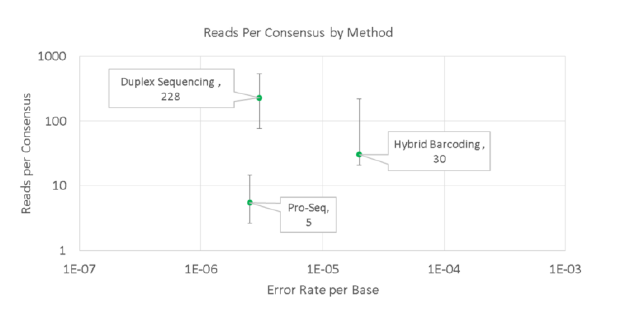
Average reads per consensus (RPC) required to represent a fixed number of input genomes, as a function of cfDNA error rate. Pro-Seq RPC was compared to estimates of RPC for duplex and hybrid barcoding using supplementary data [18]. Sequencing efficiency for Pro-Seq was calculated as on-target bases passing all Pro-Seq filters, divided by all (unfiltered) on-target bases (n = 30 runs). Error bars represent maximum and minimum reported values for the relevant data sets. Hybrid barcoding RPC estimates are also comparable to those from [24].

While measuring the molecular sensitivity and detection threshold as described above, we also observed the average background error rate (known mutants removed) for samples containing cell line DNA was 4.6 x 10^-6^ errors per base (n = 17, SD = 1.4 x 10^-6^), nearly two-fold higher and significantly different than the background rate of wild-type plasma presented above (p<0.001, t-test). This measurement is consistent with common cell line production and characterization. Cell lines are typically validated only at certain mutation positions, or if broader characterization is employed, it is typically only used on parental cell line and only with low sensitivity methods (i.e. standard NGS). Cell division, on the other hand, drives mutations in uncharacterized regions that remain undetected in cursory cell line validation, and as a result can appear as false positives or background noise in more thorough assay validation.

## Discussion

Circulating tumor DNA liquid biopsies are being evaluated, and in some cases adopted, for a number of personalized medicine applications in oncology, such as guiding treatment selection during monitoring [1, 8], minimum residual disease detection [32] and even screening (CANDACE, ClinicalTrials.gov identifier NCT02808884; CCGA, ClinicalTrials.gov identifier NCT02889978). A lower cost assay with a simple workflow and equivalent performance compared to conventional methods could increase both clinical adoption and reimbursement for these and other applications.

The measured per-base cfDNA error rate of Pro-Seq (2.6 x 10^-6^) is comparable to duplex barcoding of cfDNA (3 x 10^-6^) [18] and 10-fold better than hybrid barcoding (2 x 10^-5^) [18]. For Pro-Seq, this results in a per-base analytical specificity of 99.9997% which is better than 99.998% calculated for hybrid barcoding. Other methods [4] report similar specificity to Pro-Seq, but only for SNVs present at greater than 2% allele fraction, missing many clinically relevant mutations. The incorporation of duplex information in Pro-Seq also helps ensure that DNA damage or other early errors do not contribute significantly to background error rate. This results in extremely high per-base analytical specificity which enables detection of very low-level variants with high confidence, even on broad panels. We suspect the analytical error rate and specificity of Pro-Seq may be limited in part by real biological background, but may still improve further with implementation of *in silico* error ‘polishing’.

To the best of our knowledge, the Pro-Seq per-base detection threshold of 0.003% is among the lowest reported. Other groups have reported comparable detection thresholds when looking for multiple mutations at once [18] but this metric is not as directly reflective of assay performance. Considering the practical limits of liquid biopsy assays, we note that a detection threshold of 0.003% is safely below the maximum requirements of nearly any imaginable blood-based application. A typical human blood sample will contain on the order of a few nanograms of cfDNA per milliliter of plasma, so with a detection threshold of 0.003%, the assay technical limits are not likely to limit clinical performance in blood draws up to 100 mL volume, except in rare cases of extremely high cfDNA content per milliliter of plasma. Very low detection thresholds, however, may be important in tissue (FFPE or fresh) or other samples in cases where DNA mass is not limited and information on rare variants is desired.

Similarly, near-single-molecule sensitivity suggests that Pro-Seq is able to capture mutations present in a sample at very high efficiency, which in turn indicates that Pro-Seq does not suffer from the input template losses associated with barcoded duplex sequencing and other similar methods.

The demonstrated analytical cfDNA performance of Pro-Seq is comparable or better than conventional barcoding methods (including duplex methods), but is achieved with significantly fewer sequencing reads (∼10-fold less compared to duplex sequencing). The high reads per consensus required for duplex sequencing can at least in part be attributed to random sampling which is required to represent both senses of each starting template with sufficient redundancy to create a consensus. When sampling randomly, many other templates are sequenced unnecessarily. A less pronounced sampling effect is observed for non-duplex barcoding methods that require representation of only one sense. The sampling effect is confounded by any errors or chimeras formed within the UIDs themselves, which create isolated barcodes and requires increased sequencing [13]. Pro-Seq avoids consensus read sampling by physically linking molecules, and because no barcodes are required, avoids extra sequencing associated with barcode errors.

It should be pointed out that the sequencing redundancy for barcoding methods serves at least two functions. First, it provides the necessary number of copies to call a consensus, but additionally it provides assurance that each starting molecule is represented on the sequencer, which is required for high sensitivity applications. If every read on a sequencer was low-error, redundancy would not be required, and to minimize sequencing cost each original template would ideally be sequenced only once. However, because of sampling variation, aiming for 1x coverage of each template would result in dropout of a significant fraction of molecules, reducing assay sensitivity. Therefore, for Pro-Seq, where high accuracy individual reads are generated, a small amount of redundancy is required to ensure each starting template is represented on the sequencer. Even accounting for an extra three-fold redundancy, Pro-Seq requires comparable or fewer reads than non-duplex barcoding [24], but with better performance, and still requires more than an order of magnitude fewer reads than duplex barcoding. For even modest panel sizes, greater than 10-fold reduction in sequencing cost can result in a significant reduction in total assay cost. As panel size increases and sequencing cost becomes a larger part of the total cost, the Pro-Seq cost advantage become even more significant, on the order of 10-fold.

In addition to lower cost, Pro-Seq also provides a workflow simplicity and speed advantage, which is important for clinical adoption. In contrast to other methods which require multi-day workflows for ligation, target capture and multiple PCRs (ex. [18]), or simply multiple PCRs (ex [10]), Pro-Seq requires only a single PCR followed by cleanup, and can be completed by a single technician in a single day, with less than two hours hands on time. Also, because Pro-Seq is droplet PCR-based, it is compatible with samples containing very low DNA mass (<1ng).

The data presented in this work supports SNV and indel detection from cfDNA samples, but we expect Pro-Seq to be compatible with detection of copy number variation, loss of heterozygosity, and fusions, given appropriately designed amplicons. Pro-Seq should also find application beyond cfDNA liquid biopsy in assays that require high fidelity sequencing at low cost, such as tumor tissue sequencing and transplant monitoring. Currently, the limitations to Pro-Seq are the breadth of the assay and requirement of a droplet generation instrument. Work is ongoing to design broader panels, which we expect should be possible given the breadth achieved with other PCR assays [33]. The requirement for a droplet or emulsion generation instrument is not a significant contribution to the cost of the assay, even when the instrument cost is only amortized over a modest number of samples. Thus, labs that do not currently have droplet generation capabilities could integrate an instrument without committing to large numbers of clinical samples. Alternatively, protocols do exist for droplet/emulsion generation without the use of a dedicated instrument, and could be investigated in the future. Additionally, work is ongoing to generate a broad targeted assay using non-droplet versions of Pro-Seq (similar to S1 Fig). Even without these improvements, we expect the Pro-Seq concept to be a powerful new technology for increasing the accuracy of next generation sequencing.

Finally, the observation that wild-type cell line DNA measured in this work contains nearly two-fold higher background mutations than plasma is a powerful demonstration of Pro-Seq and an important consideration for researchers wishing to use similar reference materials in publications or assay validation. Therefore, for assay validation on low-level mutations employing cell line DNA titrations, it is important to only trust cell line DNA sequence (including wild-type cell line) at its validated positions.

## Conclusions

As described above, highly sensitive and specific circulating tumor DNA liquid biopsies have been shown to be useful in clinical applications. Error rates and clinical sensitivity continue to improve; however, clinical adoption and reimbursement remains limited, at least in part, by high assay costs. To our knowledge, the results presented here are the first to demonstrate a high performance, duplex, targeted cfDNA liquid biopsy at lower cost than conventional techniques. Pro-Seq is shown to have similar analytical sensitivity and specificity compared to gold-standard methods, but does so with reduced sequencer usage and a simple one day workflow. Additionally, Pro-Seq is able to provide duplex-based error correction, protecting against DNA damage and other spurious errors that arise from analysis of only a single strand of the template. We expect continued development of Pro-Seq to expand its breadth as well as further reduce sequencing cost, making it an attractive clinical choice for a broad range of liquid biopsy applications where low cost is an important factor.

## Acknowledgments

The authors wish to thank RainDance Technologies for droplet instrumentation and droplet reagent support.

## Author Contributions

Conceived and designed the experiments: JP, WC, AL, GS, LG, MD, LU, AM Performed the experiments: WC, AL, GS, MD, LU, LG

Analyzed the data: GS, WC, AL, LU, MD, JP Wrote the paper: JP, AM

## Materials and Methods

### DNA Isolation

cfDNA was isolated from up to 10 mL of peripheral blood per extraction. First, the blood was centrifuged for 10 min at 2,000g and 4 °C, after which plasma was removed and spun again for 10 min at 2,000g and 4 °C. cfDNA was isolated from each sample using the QIAamp Circulating Nucleic Acid Kit (Qiagen) according to the manufacturer’s instructions. DNA was eluted from the column in 0.1x IDTE (Integrated DNA Technologies (IDT)) in a two-step process. 100 µL of 0.1x IDTE was incubated in the column for 10 min, followed by a 20,000g spin for 3 min. Incubation and spin was repeated for a total elution volume of 200 µL to maximize elution yield. The full volume of DNA was further cleaned up to remove any potential inhibitors using the Monarch PCR & DNA Cleanup Kit (5 µg) (New England BioLabs (NEB)). The kit was used as per manufacturer’s instructions, except 1 mL of 2:3 binding-buffer:ethanol was added to each column in place of binding buffer alone, to improve yield. Additionally, each column was eluted in 15 µL of 0.1x TDTE. Extracted and purified DNA was then used directly for library preparation, or in cases where library preparation did not proceed within 24 hours, was frozen at -20 °C.

Following DNA extraction, the number of human genome equivalent copies in each sample was measured using quantitative PCR. Two reference loci, COG5 and ALB, were amplified in serial 10-fold dilutions and measured in duplicate.

### Targeted Pro-Seq Library Workflow

The desired number of genomic template copies for each sample was mixed into a 40 µL droplet reaction mix, containing final concentrations of 0.02 Units/µL of Q5® Hotstart DNA polymerase (NEB), 0.2 mM dNTP (NEB), 1x RDT Droplet Stabilizer (RainDance Technologies), 1x Q5® Reaction Buffer (NEB), 25 nM each gene specific Index 1 forward primer, 25 nM each gene specific Index 2 forward primer, 50 nM each gene specific reverse primer, 400 nM universal reverse primer and 200nM of the universal PEG-linked primer (S2 Fig). Nuclease-free water (IDT) was added to bring the final reaction volume to 40 µL.

Droplets were generated on the RainDrop Source instrument (RainDance Technologies) using ThunderBolts Open Source consumables (RainDance Technologies) as per manufacturer’s specifications. Approximately 8,000,0000 droplets were generated from each 40 µL sample.

Samples were then amplified on a BioRad T100 thermocycler with an initial denaturation at 98 °C for 30 s, 38 cycles at 98 °C for 10 s, 60 °C for 30 s, and 72 °C for 30 s followed by a final hold at 4 °C. Ramp rate was 1 °C/s.

Droplets were then destabilized using manufacturer’s reagents and specifications (RainDance Technologies), except 62.5 µL total destabilizer was used. Following destabilization, DNA was cleaned up using the Agencourt AMPure XP Kit (Beckman Coulter) as per manufacturer’s specification, with a 0.8:1 bead to sample ratio, and eluted in 20 µL of 0.1x IDTE.

Un-linked DNA was then digested enzymatically in a 50 µL reaction by mixing the eluted DNA from the previous step with 6.7 Units of T7 Exonuclease (NEB), 41.7 units of RecJf (NEB) and nuclease free water (IDT) in a 1x final concentration of NEBuffer 4 (NEB). Digestion proceeded at 37 °C for 1 h, followed by a 70 °C inactivation step for 20 min. Following digestion, DNA was cleaned up using the Agencourt AMPure XP Kit (Beckman Coulter) as per manufacturer’s specification, with a 1.6:1 bead to sample ratio, and eluted in 25 µL of 0.1x IDTE. After cleanup, the sample was ready for sequencing. Standard amplicon libraries were generated in the same way, but with an un-linked version of the universal PEG-linked primer, and no digestion step.

### Pro-Seq Panels

The Pro-Seq ten-amplicon panel covers the regions described in S4 Table. The seven-amplicon panel is the same but does not include TP53, GNAS or EGFR exon 19.

### DNA Sequencing

All sequencing for this work was performed on a MiSeq (Illumina), though Pro-Seq has also been demonstrated on the two-color MiniSeq platform (Illumina). Prior to sequencing, samples were quantified using the KAPA Library Quant Kit (KAPA Biosystems). Samples were loaded onto the MiSeq with a modified protocol as follows: 18 µL of library was mixed with 2 µL of 1 N NaOH and incubated for 5 min at room temperature, and then placed on ice. The sample was then mixed with denatured PhiX (to 5% of library concentration), 2 µL of 1 N HCl and diluted to 600 µL with Illumina HT1 buffer (final library concentration is 5.5 pM). The resulting mix was then loaded onto the sequencer as per the manufacturer’s protocol.

Custom Read 1, Index 1 and Index 2 primers were used and loaded as per manufacturer’s instructions. Read 1 was a 1:1 mix of each of Index 1 and Index 2 primers, in addition to the standard Read 1 primer. Each index read used a custom index primer along with the standard Read 1 primer. A Custom Read 2 primer was also used by adding 3.5 µL of 100 µM custom Read 2 primer into the Read 2 Primer well (MiSeq cartridge well 14). The length of Read 1 and Read 2 were configured to overlap for each amplicon, specified in the sequencer sample sheet. Custom Index 1 and Index 2 reads were utilized to verify the presence of both starting strands in each analyzed cluster (S2 Fig). To do this, the ‘2Read2Index.xml’ file on the sequencer was modified to perform the Index 2 read before sequencing turnaround, and subsequently omit the dark cycles. The ‘Reads.xml’ file was modified to sequence the correct number of cycles for the Index 2 read, and the ‘Chemistry.xml’ file was modified to support the Index 2 read before turnaround.To enable our custom analysis, additional data beyond the FASTQ files was collected. ‘Configuration.xml’ was modified to save intensity files for each cycle, and ‘MiSeq Reporter.exe.config’ was modified to keep reads that did not pass filter, and generate FASTQ files for the index reads. A noteworthy advantage of Pro-Seq is that useful data is collected even from clusters that do not pass Illumina’s filtering schemes, further improving efficiency over other methods.

### Data Processing

Due to the unique nature of Pro-Seq, custom scripts were written for both SNV and indel analysis. The general analysis pipeline is outlined in S3 Fig. For SNVs, BWA-MEM was used to filter all unaligned or malformed clusters. Clusters were discarded if they did not align to the panel or the alignments did not make sense in reference to the genome. After alignment, a custom second filtering was applied for doubly-seeded (DS) clusters, eliminating clusters that only represented one of the two expected index reads (DS clusters contained the expected sequence in both index reads). Next, each doubly-seeded cluster was analyzed for the presence of mixed signal, which indicates an error that was not automatically corrected during sequencing. Mixed bases (not entire reads or clusters) were identified and masked by comparing the relative fluorescent intensities (fQ) of each nucleotide for a given cycle, read and cluster, as well as the quality score at that position. Next, sequencing reads were compiled per cluster to determine a base call for each reference position on the panel, taking masked bases into account. Typically, at least two base calls per position must agree for a base call to be made for a given cluster. After base calls were made for each cluster, base calls for each amplicon on the panel were compiled and non-reference calls identified by our custom variant caller, typically set to call SNVs above two genome equivalent copies present in the original starting sample. A simpler custom analysis was employed for indels, since Pro-Seq is not typically required for their detection. Read 1 and Read 2 are merged within each cluster, aligned with BWA-MEM, and malformed or off-panel clusters are discarded. Primer regions are then trimmed, and inter-primer regions are grouped based on indels. Potential indels are then fed to the variant caller which checks for sequencing artifacts against an internal library, and typically calls valid indels above two genome equivalent copies.

### Molecular Sensitivity and Specificity

Molecular sensitivity was measured by titrating characterized mutant cell line DNA into wild-type plasma DNA. First, 1% mutant cell line DNA (Multiplex 1 cfDNA Reference HD778, Horizon Discovery, Cambridge, UK) was measured for mutant content using the Pro-Seq ten-amplicon panel to verify the manufacturer’s reported allelic frequency as shown in S5 Table. Excellent concordance was observed between expected and measured values for all five panel mutants. Mixtures of 1% cell line DNA and wild-type plasma DNA (IPLAS – K2 EDTA, Innovative Research, Novi, MI) samples were created with 0.3%, 0.1%, 0.03%, 0.01% and 0.00% average individual allelic frequency. 15,000 genome equivalents were analyzed in each sample, resulting in 45, 15, 4.5, 1.5 and 0 average mutant copies, respectively. Each sample was run in duplicate, for a total of ten mutants measured per dilution. Mutants were called positive if present at >0.5 genome equivalent copies.

Detection threshold was measured by creating samples with 1000 genome equivalent copies of 1% mutant cell line DNA (Multiplex 1 cfDNA Reference HD778, Horizon Discovery, Cambridge, UK), so that each of the five mutants shown in S5 Table were nominally present at ten copies each (above the molecular sensitivity limit). Additional wild-type cell line DNA (Custom cfDNA Reference HD-C328, Horizon Discovery, Cambridge, UK), with mutants shown in S5 Table specified to be present at 0.00% by the manufacturer, was added to the 1% cell line DNA to reach the desired mutation frequencies (S6 Table). Each sample was run in duplicate, for a total of ten mutants measured per dilution. Wild-type only controls were also run at the highest input mass to detect any low-level mutant background signal. Measured mutation frequency for each sample is shown in S6 Table.

Cell line DNA was used in place of plasma-derived cfDNA in these experiments due to the large DNA mass required to meet the low detection thresholds (∼1 µg for each the 0.003% samples).

## Supporting Information

**Fig S1.**
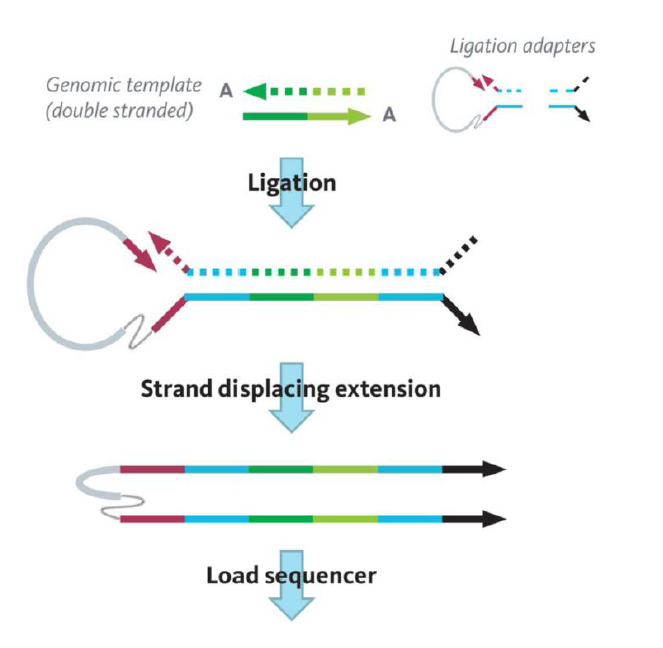
Overview of Whole Genome Pro-Seq. In brief, double-stranded DNA is ligated with unique PEG-linked ‘loop adapters’. The bound priming site on the loop adapter undergoes a single extension with a strand displacing polymerase to generate a molecular construct where each construct and Pro-Seq cluster contains representation from each sense of the starting molecule. After cleanup and quantification, the library is ready to load on the sequencer. This PCR-free workflow is very rapid and can be performed by a single technician in less than four hours (less than two hours hands-on time).

**Fig S2.**
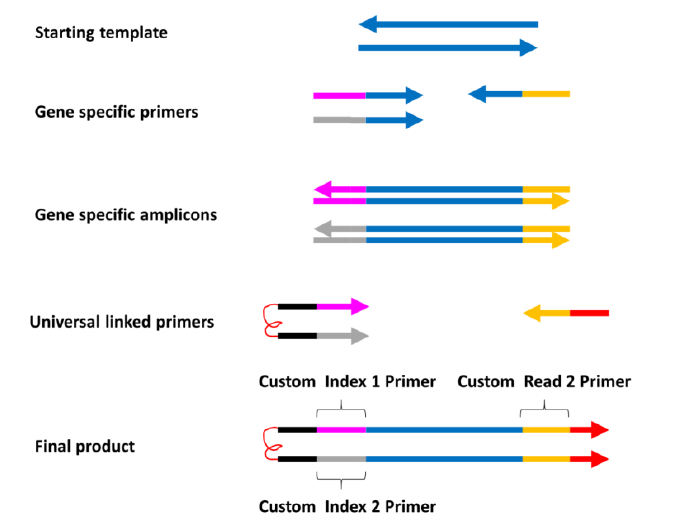
Pro-Seq PCR and sequencing architecture. On average, zero or one DNA templates were loaded into each droplet, along with other background DNA (DNA that is not amplified by gene specific primers). Each droplet also contained multiplexed gene specific primers, and universal linked primers. In this work, between seven and 19 amplicons were multiplexed together. Each amplicon used two gene specific forward primers with different linking sequences (pink, grey) to the universal linked primer, which enabled identification of Pro-Seq clusters on the sequencer, along with a single gene specific reverse primer. The two different forward gene specific primers per amplicon created two gene specific amplicon types per target, such that when two linker primers were used, on average both senses of the starting templates were represented in 50% of the Pro-Seq clusters (as the number of linker primers increases, the fraction of clusters representing both senses also increases). Universal 5’ PEG-linked primers containing flow cell adapter sequences (black) extended off the two gene specific amplicons with a single universal reverse primer that contained the second flow cell adapter sequence (red). After sufficient cycling, all universal linkers were ‘filled’ to create the final sequenced product. Not shown is the un-linked reverse complement of the final product which was digested after emulsion breaking, prior to sequencing. Sequencing primer locations were as indicated.

**Fig S3.**
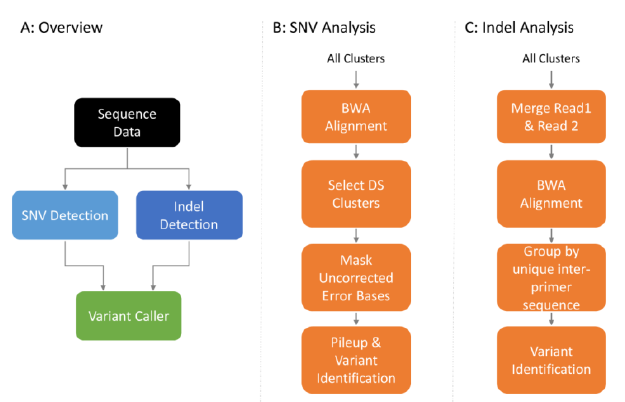
Pro-Seq analysis pipeline. (A) Full analysis overview. SNV and indel detection were handled separately, after which a combined variant caller identified any non-reference sequences. (B) SNV analysis consisted of alignment, doubly-seeded (DS) cluster selection, error base masking (to eliminate remaining errors not corrected during sequencing) and then pileup and variant identification. (C) Indel analysis consisted of alignment, trimming of known primer sequences and grouping by specific inter-primer sequences. Inter-primer sequences were piled up, followed by variant identification.

**S4 Table:** In brief, double-stranded DNA is ligated with unique PEG-linked ‘loop adapters’. The bound priming site on the loop adapter undergoes a single extension with a strand displacing polymerase to generate a molecular construct where each construct and Pro-Seq cluster contains representation from each sense of the starting molecule. After cleanup and quantification, the library is ready to load on the sequencer. This PCR-free workflow is very rapid and can be performed by a single technician in less than four hours (less than two hours hands-on time).

| Amplicon | Gene | Exon | Chromosome | GRCh38 start position | GRCh38 stop position |
| --- | --- | --- | --- | --- | --- |
| 1 | ALB | 12 | 4 | 73418216 | 73418313 |
| 2 | BRAF | 15 | 7 | 140753297 | 140753396 |
| 3 | COG5 | 12 | 7 | 107298099 | 107298192 |
| 4 | EGFR | 19 | 7 | 55174728 | 55174812 |
| 5 | EGFR | 20 | 7 | 55181353 | 55181415 |
| 6 | EGFR | 21 | 7 | 55191768 | 55191864 |
| 7 | KRAS | 2 | 12 | 25245310 | 25245395 |
| 8 | GNAS | 8 | 20 | 58909321 | 58909432 |
| 9 | PIK3CA | 10 | 3 | 179218238 | 179218343 |
| 10 | TP53 | 8 | 17 | 7673781 | 7673869 |

**S5 Table:** Measured vs. expected allelic frequency for the five cell line DNA mutants that are contained within the ten-amplicon Pro-Seq panel.

| Gene | Variant | Expected Allelic Frequency | Measured Allelic Frequency |
| --- | --- | --- | --- |
| EGFR | T790M | 1.0% | 0.78% |
| EGFR | L858R | 1.0% | 0.98% |
| KRAS | G12D | 1.3% | 1.1% |
| PIK3CA | E545K | 1.3% | 1.1% |
| EGFR | ΔE746 - A750 | 1.0% | 0.91% |

**S6 Table. Expected vs. measured allelic frequency for the five cell line DNA mutants used in the detection threshold measurements.** The second replicate of 0.1% had low EGFR L858R representation compared to other mutants, but still above one template copy, and may be due to sampling variation.

